# Live applications of norbormide-based fluorescent probes in *Drosophila melanogaster*

**DOI:** 10.1101/517334

**Authors:** Alessia Forgiarini, Zifei Wang, Claudio D’Amore, Morgan Jay-Smith, Freda Fan Li, Brian Hopkins, Margaret A. Brimble, Andrea Pagetta, Sara Bersani, Sara De Martin, Barbara Napoli, Sergio Bova, David Rennison, Genny Orso

**Affiliations:** Department of Pharmaceutical and Pharmacological Sciences, University of Padova, Padova, Italy; University of Auckland, School of Chemical Sciences, Auckland, NZ; Landcare Research, Lincoln, New Zealand; Scientific Institute IRCCS Eugenio Medea, Bosisio Parini, Lecco, Italy

## Abstract

In this study we investigated the performance of two norbormide (NRB)-derived fluorescent probes, NRB^MC009^ (green) and NRB^ZLW0047^ (red), on dissected, living larvae of *Drosophila*, to verify their potential application in confocal microscopy imaging *in vivo*. To this end, larval tissues were exposed to NRB probes alone or in combination with other commercial dyes or GFP-tagged protein markers. Both probes were rapidly internalized by most tissues (except the central nervous system) allowing each organ in the microscope field to be readily distinguished at low magnification. At the cellular level, the probes showed a very similar distribution (except for fat bodies), defined by loss of signal in the nucleus and plasma membrane, and a preferential localization to endoplasmic reticulum (ER) and mitochondria. They also recognized ER and mitochondrial phenotypes in the skeletal muscles of fruit fly models that had loss of function mutations in the atlastin and mitofusin genes, suggesting NRB^MC009^ and NRB^ZLW0047^ as potentially useful *in vivo* screening tools for characterizing ER and mitochondria morphological alterations. Feeding of larvae and adult *Drosophilae* with the NRB-derived dyes led to staining of the gut and its epithelial cells, revealing a potential role in food intake assays. In addition, when flies were exposed to either dye over their entire life cycle no apparent functional or morphological abnormalities were detected. Rapid internalization, a bright signal, a compatibility with other available fluorescent probes and GFP-tagged protein markers, and a lack of toxicity make NRB^ZLW0047^ and, particularly, NRB^MC009^ one of the most highly performing fluorescent probes available for in vivo microscopy studies and food intake assay in *Drosophila*.

## Introduction

Norbormide [5-(α-hydroxy-α-2-pyridylbenzyl)-7-(α-2-pyridylbenzylidene)-5-norbornene-2,3-dicarboximide] (NRB) is a selective rat toxicant that exhibits little or no non-target effects (1), and was developed and commercialized as an ecologic pesticide in the 1980s(Roszokwski, 1965). Evidence suggests that the rat-selective action of NRB is mediated by a generalized vasoconstrictor effect that has only been observed in the rat peripheral blood vessels, both *in vivo* and *in vitro*. In contrast, NRB displays a vasorelaxant action in arteries from non-rat species, as well as in rat aorta and extravascular smooth muscle, that has been proposed to be the result of a reduction of Ca^2+^ entry through L-type Ca^2+^ channels (2,3). The molecular mechanism underlying NRB-induced vasoconstriction is not known, however, it has been proposed that the compound acts on rat vascular myocytes where it activates the PLC-IP3-PKC pathway (2), a signaling cascade stimulated by most receptor-coupled vasoconstrictor agents (4). In an attempt to identify the cellular targets of NRB, we previously developed fluorescent derivatives of the parent compound by linking it to either nitrobenzoxadiazole (NBD) or boron-dipyrromethene (BODIPY FL) fluorophores, and found that both were able to clearly label intracellular structures such as endoplasmic reticulum (ER), Golgi apparatus, mitochondria, and lipid droplets (LDs) in various cell lines, in the absence of cytotoxic effects. Based on these results, we proposed NRB as a scaffold for the development of new, high performing, non-toxic fluorescent probes for live cell imaging (5,6).

*Drosophila melanogaster* is an animal model widely used to investigate the biochemical pathways and the cellular/subcellular morphological alterations that characterize human diseases (Aldaz et al., 2010; Chatterjee, 2014; Mushtaq et al., 2016; Musselman and Kühnlein, 2018; Orso et al., 2005; Tan and Azzam, 2017) Confocal fluorescent microscopy live imaging is a particularly informative methodology in this model, especially when fluorescent probes are used in combination with genetic tools (e.g. mutant flies, RNA interference, fluorescently marked proteins, Gal4/UAS activation system) (13,14), allowing the visualization of dynamic biological processes in living systems without the artifacts often generated by sample fixation procedures (15).

In this study, we investigated the performance of two NRB-derived fluorescent probes, the previously developed NRB^MC009^ (green fluorescence) and the newly developed NRB^ZLW0047^ (red fluorescence), on dissected, living third instar larvae of *Drosophila melanogaster*, to assess their potential application in confocal microscopy imaging *in vivo*. In particular, we were able to characterize the distribution of NRB^MC009^ and NRB^ZLW0047^ to cellular structures and organelles in tissues of wild-type *Drosophila* larvae, as well as in tissues of mutant lines exhibiting morphological alterations of endoplasmic reticulum and mitochondria. Finally, we explored if these probes could be useful for studying fruit fly feeding and gut morphology.

## Material and methods

### NRB^MC009^ and NRB^ZLW0047^ synthesis

NRB^MC009^, a BODIPY FL derivative of norbormide, was synthesized as previously reported (6). Stock solutions 1 mM in DMSO were prepared and maintained at −20 °C and diluted to the desired concentration before each experiment.

NRB^ZLW0047^, a BODIPY TMR derivative of norbormide, was prepared as follows. BODIPY TMR (4,4-difluoro-5-(4-methoxyphenyl)-1,3-dimethyl-4-bora-3a,4a-diaza-s-indacene-2-propionic acid) (16), along with its corresponding N-hydroxysuccinimide ester, BODIPY TMR NHS ester (16), and N-2’-aminoethyl-endo-5-(α-hydroxy-α-2-pyridylbenzyl)-7-(α-2-pyridylbenzylidene)-5-norbornene-2,3-dicarboximide (17) were prepared using literature methods. A solution of N-2’-aminoethyl-endo-5-(α-hydroxy-α-2-pyridylbenzyl)-7-(α-2-pyridylbenzylidene)-5-norbornene-2,3-dicarboximide (111 mg, 0.20 mmol), BODIPY TMR NHS ester (110 mg, 0.20 mmol) and *N,N*-diisopropylethylamine (35 µl, 0.20 mmol) in dichloromethane (7 ml) was stirred at room temperature for 16 h. The mixture was then diluted with dichloromethane (20 ml), washed with water (10 ml), the separated aqueous phase further extracted with dichloromethane (2 × 10 ml), the combined organic layers washed with brine (3 × 20 ml), dried over anhydrous magnesium sulfate, filtered and the solvent removed in vacuo. Purification by flash chromatography (petroleum ether/ethyl acetate, 1:3) afforded NRB^ZLW0047^ as a mixture of endo stereoisomers (purple solid; 70 mg, 37%). ^1^H NMR (400 MHz, CDCl_3_) *δ* 8.64-8.38 (2H, m, αPyr), 7.88-7.82 (2H, m, ArH), 7.59-6.77 (20H, m, ArH), 6.53-6.47 (1H, m, C=CH), 6.29 (0.2H, br s, OH), 6.28 (0.1H, br s, OH), 6.21-6.20 (0.2H, m, W/H-6), 6.15-6.14 (0.1H, m, U/H-6), 5.54-5.53 (0.4H, m, Y/H-6), 5.52-5.51 (0.3H, m, V/H-6), 5.22 (0.4H, br s, OH), 5.15 (0.3H, br s, OH), 4.49-4.46 (0.2H, m, W/H-1), 4.46-4.43 (0.4H, m, Y/H-1), 4.31-4.29 (0.3H, m, V/H-1), 4.03-4.01 (0.1H, m, U/H-1), 3.85-3.84 (3H, m, OMe), 3.84-3.24 (7H, m, H-2, H-3, H-4 and NCH_2_CH_2_), 2.75-2.72 (2H, m, COC*H*_2_CH_2_ or COCH_2_C*H*_2_), 2.47 (3H, m, Me), 2.26-2.22 (2H, m, COC*H*_2_CH_2_ or COCH_2_C*H*_2_), 2.20-2.18 (3H, m, Me).

Stock solutions 1 mM in DMSO were prepared and maintained at −20 °C and diluted to the desired concentration before each experiment.

### Fluorescence spectra

Excitation and emission spectra of NRB^MC009^ and NRB^ZLW0047^ were obtained by diluting the stock solution in Milli-Q water to reach the final concentration of 2 μM. To verify that the medium used for *Drosophila* live imaging (HL3) had no effect on the emission spectra, a comparison was made between probes diluted in water and probes diluted in HL3 l - no variation in fluorescence was observed.

Analysis of excitation/emission peaks were evaluated using a Jasco FP6500 spectrofluorometer (temperature 25 °C; b=1 cm; λ ex/em: 470/540 for NRB^MC009^ and 545/580 for NRB^zlw0047^; sli: 5/10 nm; data pitch 0.2 nm; scanning speed 200nm/min).

### Fly stocks

*Drosophila melanogaster* strains used: w^[1118]^ (BL-5905), Tubulin-Gal4 (BL-5138), UAS-Mito-GFP (BL-8443), UAS-mCD8-GFP (BL-5130), were obtained from Bloomington Drosophila Stock Center, and UAS**-**ATL2^RNAi^ (18) and UAS-Marf^RNAi^ (ID 40478), were resourced from Vienna Drosophila Resource Center. UAS-Lamp-GFP was provided by Helmut Krämer (University of Texas Southwestern Medical Center, Dallas), and UAS-HneuGFP was generated by cloning HNEU-GFP (19) in pUASTattB, and transgenic lines generated by BestGene Inc, (Chino Hills, CA, USA). W^[1118]^ flies were maintained on standard food at 25 °C, and Gal4/UAS crossings were performed at 28 °C. Starvation was induced by leaving third instar larvae for 6 hours in 20% sucrose dissolved in PBS.

### Larval dissection

Fly larvae having reached the third instar stage were pinned between the posterior spiracles and above the mouth hooks in a Sylgard dissection dish, and cut along the dorsal midline. Hemolymph-like (HL3) saline (70 mM NaCl, 5 mM KCl, 1.5 mM CaCl_2_, 20 mM MgCl_2_, 10 mM NaHCO_3_, 5 mM trehalose, 115 mM sucrose, 5 mM sodium HEPES, pH 7.2, all supplied by Sigma-Aldrich) was added and the lateral flaps were fastened with four needles to stretch the body wall. All larval organs were left on the muscle fillet for whole larval acquisition, whereas for single tissue acquisition the unnecessary organs were removed.

### Live tissue imaging

To characterize NRB^MC009^ and NRB^ZLW0047^ localization, each were diluted in HL3 medium at concentrations of 500 nM and 1 µM, respectively, and were added to the dissected larva and images acquired after 15 min. For NRB^MC009^ colocalization studies, ER-Tracker™ Red 2 µM (BODIPY™ TR Glibenclamide), MitoTracker™ Orange CMTMRos 1 µM, LysoTracker™ Deep Red 2 µM, HCS LipidTOX™ Deep Red Neutral Lipid Stain 1:100, or CellMask 1 µM (all by Thermo Fisher, respectively #E34250, #M7510, #L12492, #H34477, and #10045) were added, together with NRB^MC009^ at 500 nM. To verify NRB^ZLW0047^ colocalization with other organelles, it was added on dissected larvae expressing GFP-tagged proteins (Hneu-GFP, Mito-GFP, Lamp-GFP, and mCD8-GFP), or together with BODIPY 493/503™ dye 10 μg/ml (#D3922, Thermo Fisher). Whole larvae images and magnifications were acquired using a Zeiss LSM800 Axio Observer Z1 inverted microscope equipped with a Zeiss Plan-Apochromat 5x/0.15 ph1 or 40x/0.95 objectives, all other images were acquired with a Nikon EZ-C1 confocal microscope equipped with a Nikon Plan Apo 60×/1.40 or a Nikon Plan Apo 40x/1.0 oil immersion objectives.

### Three-choice preference assay

To test larval food preference, a Petri dish was divided in three quadrants filled with a warm liquefied standard food solution. In two quadrants, the food contained either NRB^MC009^ or NRB^ZLW0047^ (both 20 µM), in the third the food was left probe-free. At the beginning of the experiment, a group of 10 larvae was placed in the middle of the assay plate and after 5 min the number of larvae on each quadrant was counted (20). The experiment was repeated five times.

### Food intake test

The food intake test was conducted in third instar larvae starved for 1 hour in 20% sucrose dissolved in PBS. The starved larvae were then divided into three groups, separately fed with brilliant blue R dye 0.08%, NRB^MC009^, or NRB^ZLW0047^ (both at a concentration of 20 µM) dissolved in liquid food (sucrose 20% and 20% dry yeast in PBS) for 30 min (21), frozen at −80° C and imaged using a Leica MZ 16 FA microscope. To estimate food intake, the area of dye-labeled gut, visible through the cuticle, was quantified relative to larval total body area. 10 larvae for each experiment were used and the experiment was repeated three times. Image analysis was performed with ImageJ 1.52h software.

### Analysis of gut fluorescence in *Drosophila* chronically fed with NRB^MC009^ and NRB^ZLW0047^

Third instar larvae, grown in NRB^MC009^- and NRB^ZLW0047^-enriched food as described in the chronic toxicity assay, were collected and food debris was removed by washing in PBS for 5 minutes and 70% ethanol for 1 minute and then dissected, being careful to not cut the digestive tract. Images were acquired using a Zeiss Axio Observer Z1 inverted microscope equipped with a Zeiss Plan-Apochromat 5x/0.15 ph1 or 40x/0.95 objectives.

### Chronic toxicity assay

The toxicity of NRB-derived probes was investigated by exposing the flies to NRB^MC009^ or NRB^ZLW0047^ 20 µM over the entire life-cycle (mating, eggs maturation, pupal development, eclosion). Male and female w^[1118]^ flies were placed in a tube with standard food containing vehicle, NRB^MC009^, or NRB^ZLW0047^ 20 µM, and grown at 25 °C. After 5-6 days, parent flies were removed and analyzed to verify probe intake. Eggs were left to develop in food containing one of each of the NRB fluorescent probes until the eclosion. Two parameters were considered in the evaluation of toxicity: eclosion rate (percent of emerged flies versus the total number of pupae, including the dead ones); morphological alterations of eclosed adults. Morphological alterations of male and female adult body, eyes, wings, and legs were evaluated and imaged using a Leica MZ 16 FA microscope.

### Statistical analysis

Analysis of colocalization was performed using Pearson’s correlation coefficient calculated with Coloc2 plugin of Fiji (22). All values are expressed as means ± standard error of the mean (SEM) of n observations (n≥10). Significance was calculated using One-way ANOVA test followed by Tukey’s Multiple Comparison Test, using GraphPad Prism 3.03.

## Results and discussion

### Synthesis and excitation/emission spectra of fluorescent probes NRB^MC009^ and NRB^ZLW0047^

The structures and synthetic routes to generate NRB^MC009^ and NRB^ZLW0049^ are summarized in Figs 1A and 1B, respectively. The excitation and emission spectra of the dyes in water, and in the physiological buffer (HL3) solution which was used to perform live imaging experiments in *Drosophila* tissues, are presented in Figs 1C and 1D, respectively, and summarized in Fig 1E. The excitation spectra analysis showed that the probes can be excited using the common laser lines 561 nm (NRB^ZLW0049^) and 488 nm (NRB^MC009^), respectively.

**Fig 1.**
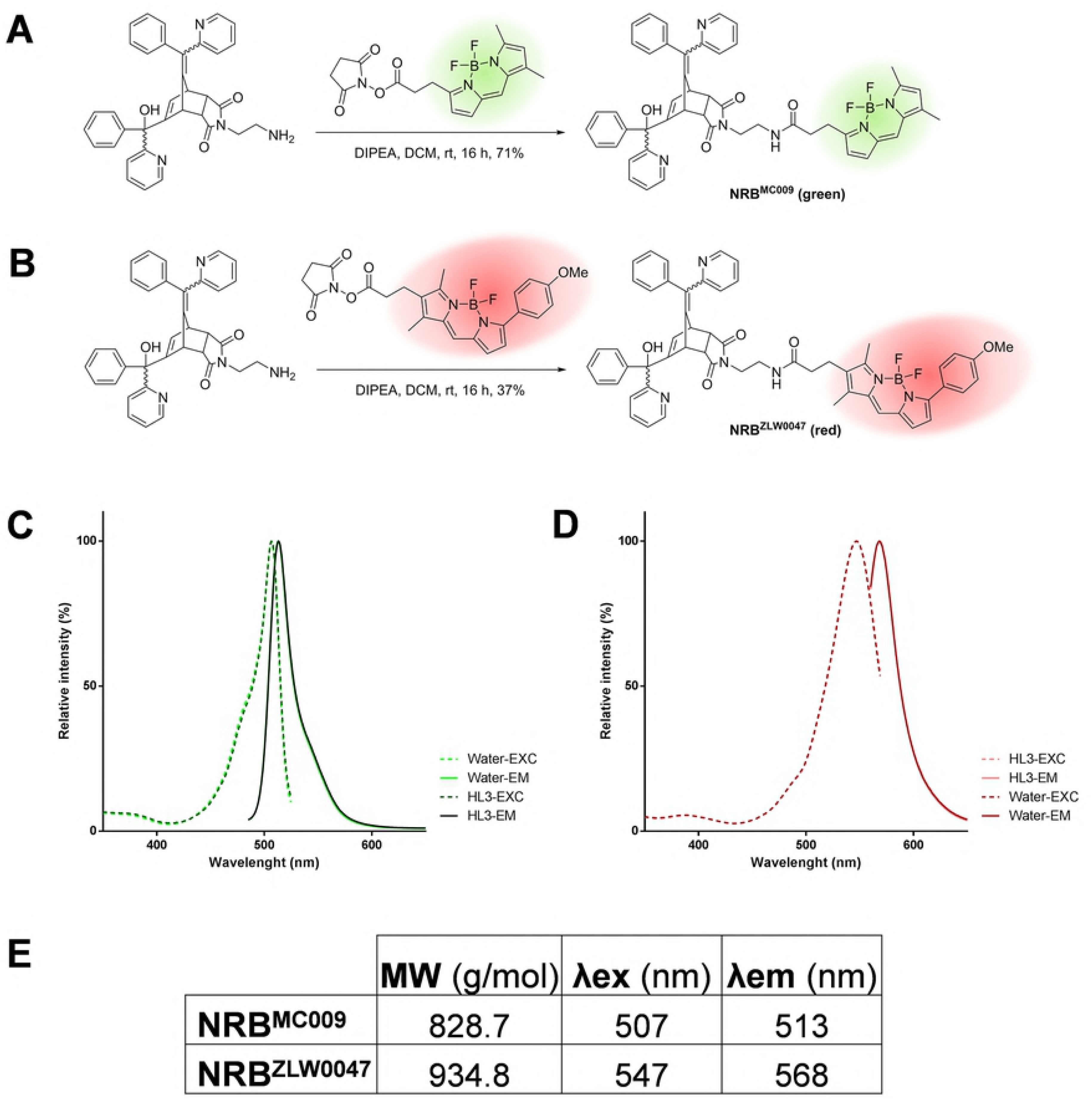
NRB^MC009^ and NRB^ZLW0047^ synthesis and fluorescent spectra. Scheme of the synthesis of NRB^MC009^ (**A**) and NRB^ZLW0047^ (**B**). Excitation and emission spectra of NRB^MC009^ (**C**) and NRB^ZLW0047^ (**D**) diluted in water (light color) or in HL3 medium (dark color). Summary table of physical properties of NRB^MC009^ and NRB^ZLW0047^ (**E**), MW: molecular weight; λex: excitation wavelength; λem: emission wavelength.

### Tissue distribution of NRB^MC009^ and NRB^ZLW0047^ in *Drosophila* larvae

To characterize NRB^MC009^ and NRB^ZLW0047^ fluorescence distribution in *Drosophila* tissues, we dissected a wild type larva and exposed the whole body to the fluorescent probes. Figs 2A and 3A are confocal images of dissected third instar larvae in which all tissues were left intact and labeled with 500 nM NRB^MC009^ or 1 μM NRB^ZLW0047^, respectively. As shown in the pictures, both dyes localized to most of the tissues, with an apparent brighter signal detected in imaginal discs, tracheal system, salivary glands, and fat body (Figs 2B, D, H, J, respectively, and Figs 3B, D, H, J, respectively), and a less intense signal being detected in muscular tissues, oenocytes, the entire digestive tract, the ring gland, and epidermal cells (Figs 2C, E, F, G, I, K, respectively, and Figs 3C, E, F, G, I, K, respectively). NRB^MC009^ and NRB^ZLW0047^ were unable to label the central nervous system i.e. ganglion, brain lobes, and nerves (dotted boxes on Figs 2A and 3A).

**Fig 2.**
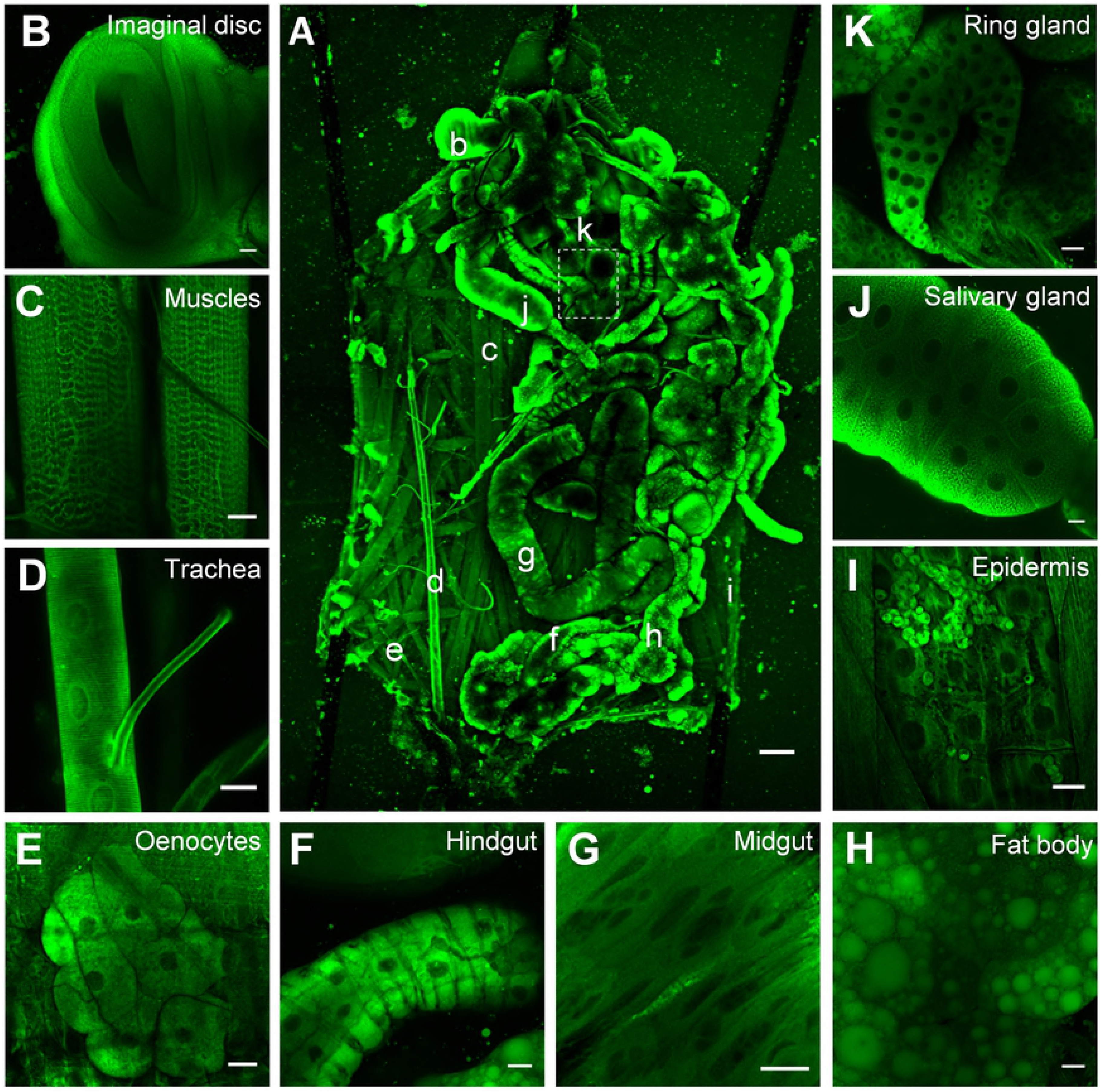
NRB^MC009^ distribution in *Drosophila* larval tissues. Confocal live imaging of (**A**) whole dissected w^[1118]^ third instar larva labeled with NRB^MC009^ 500 nM. Small letters reveal corresponding magnified tissues, dotted box shows unlabeled CNS. Magnification 5x; scale bar 200 μm. Detailed images of (**B**) leg imaginal disc, (**C**) muscles, (**D**) trachea, (**E**) oenocytes, (**F**) hindgut, (**G**) midgut, (**H**) fat body, (**I**) epidermis and hematocytes, (**J**) salivary gland, and (**K**) ring gland; magnification 40x; scale bars 20 μm.

**Fig 3.**
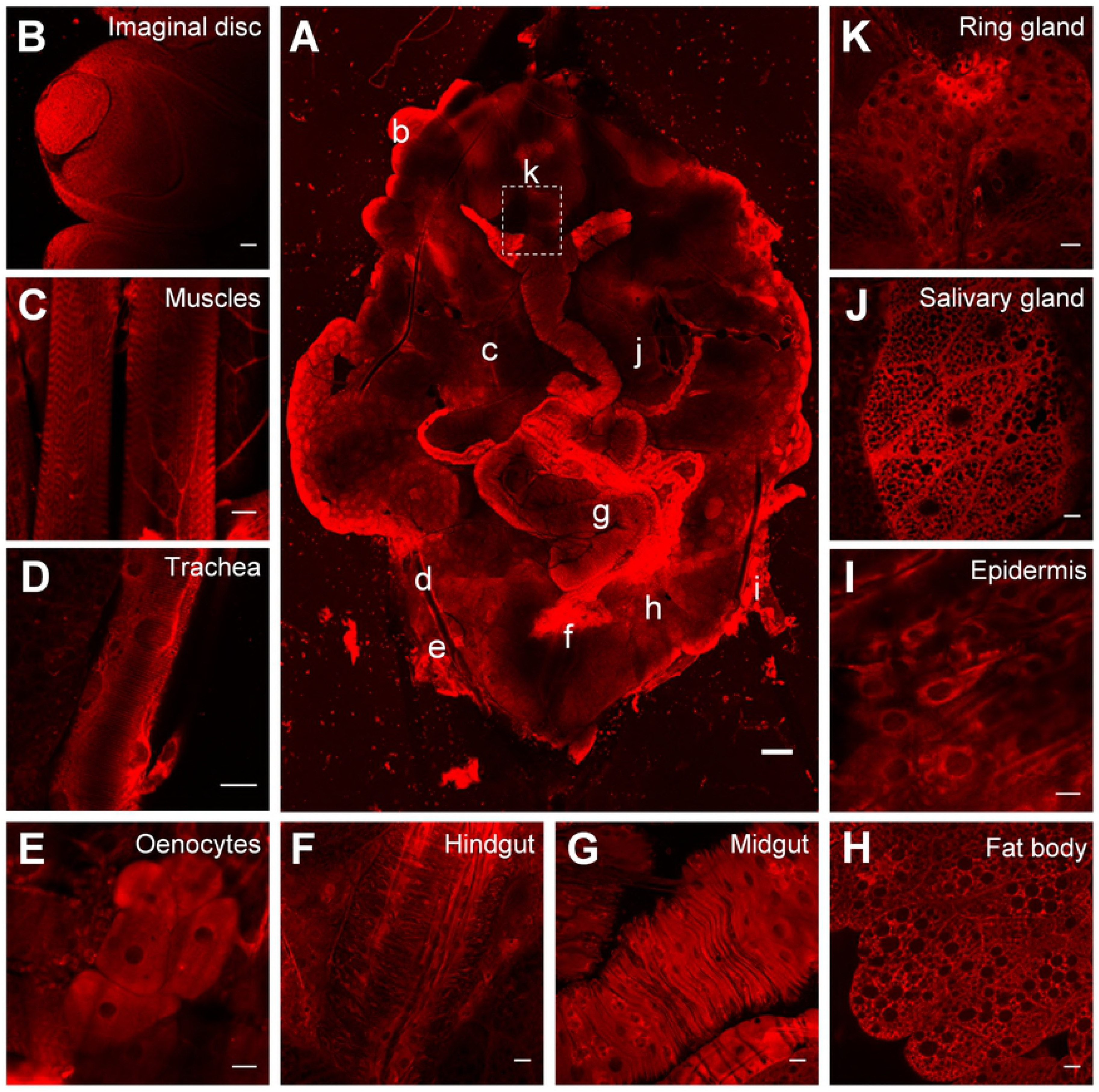
NRB^ZLW0047^ distribution in *Drosophila* larval tissues. Confocal live imaging of (**A**) whole dissected w^[1118]^ third instar larva labeled with NRB^ZLW0047^ 1 µM. Small letters reveal corresponding magnified tissues, dotted box shows unlabeled CNS. Magnification 5x; scale bar 200 μm. Detailed images of (**B**) leg imaginal disc, (**C**) muscles, (**D**) trachea, (**E**) oenocytes, (**F**) hindgut, (**G**) midgut, (**H**) fat body, (**I**) epidermis, (**J**) salivary gland, and (**K**) ring gland; magnification 40x; scale bars 20 μm.

The labeling properties of the two probes were compared by co-loading larval tissues with both dyes, and merging the corresponding images for colocalization analysis. The results, reported in Fig 4A-D, indicate that both dyes were able to penetrate the cells of the tissues, and effectively label the intracellular structures. Cell internalization of the dyes was very rapid, allowing a clear visualization of the intracellular structures in less than 1 min (data not shown). Pearson’s coefficient results (Figs 4E-F) confirmed that NRB^MC009^ and NRB^ZLW0047^ recognized the same intracellular structures in most of the tissues investigated. However, a clear difference in subcellular expression between NRB^MC009^ and NRB^ZLW0047^ was observed in the fat bodies, in which lipid droplets (LDs) were selectively stained by NRB^MC009^ (Fig 4B). This behavior may reflect a different binding capacity of the two probes to the constituents of LDs, i.e. neutral lipids, mainly triacylglycerols and sterol esters, and phospholipids (23,24).

**Fig 4.**
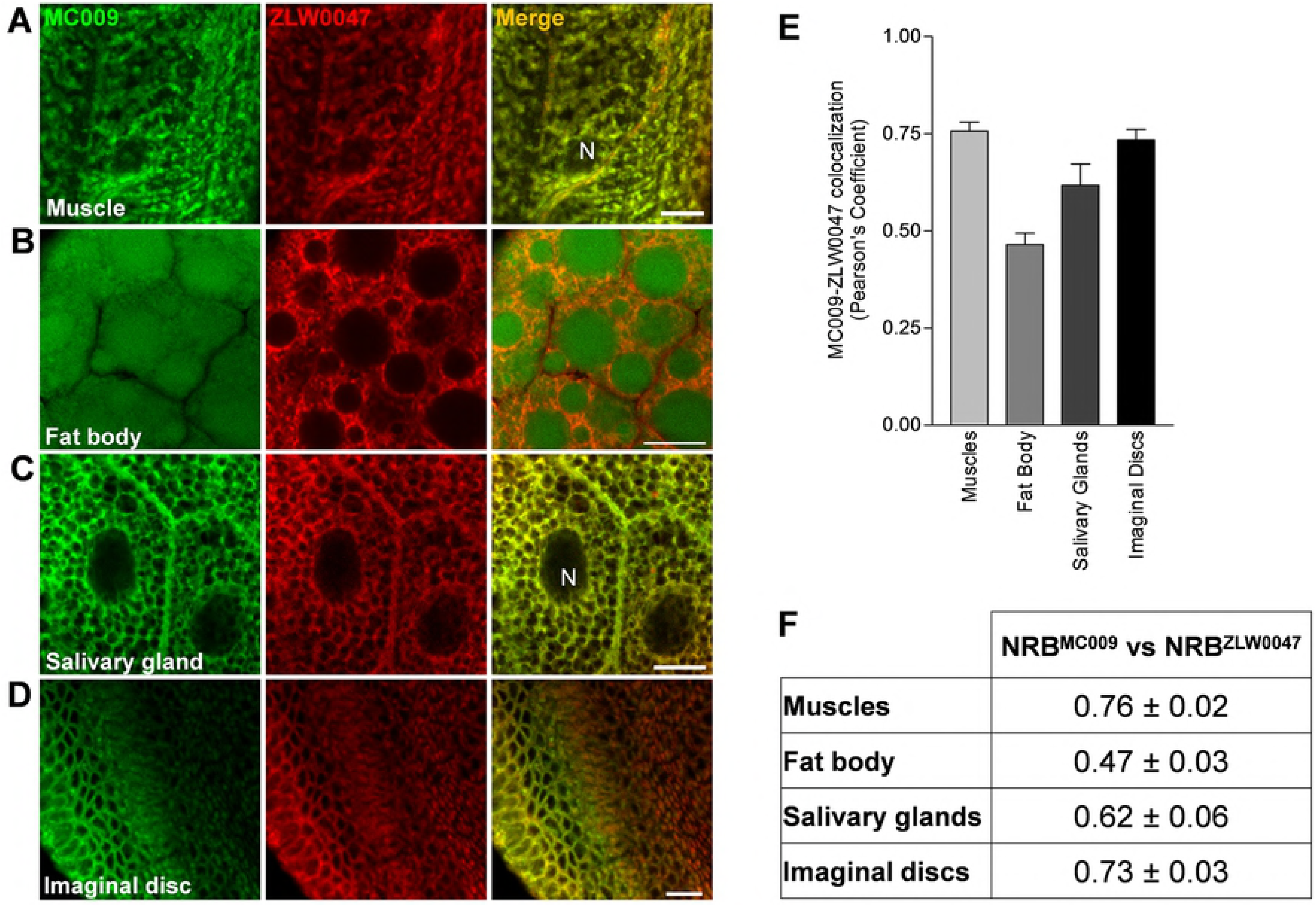
NRB^MC009^ and NRB^ZLW0047^ colocalization. Confocal live imaging of dissected w^[1118]^ third instar larval (**A**) muscle, (**B**) fat body, (**C**) salivary gland, and (**D**) imaginal disc labeled with NRB^MC009^ 500 nM (green) together with NRB^ZLW0047^ 1 µM (red). Magnification 60x; scale bars 10 μm. Graph (**E**) and summary table (**F**) of Pearson’s correlation coefficients between NRB^MC009^ and NRB^ZLW0047^ in the evaluated tissues. Data expressed as mean ± SEM, n≥10.

### Cellular distribution of NRB^MC009^ and NRB^ZLW0047^

We next characterized the cellular structures labeled by NRB^MC009^ and NRB^ZLW0047^. In all the tissues investigated, we found that both probes allowed the visualization of intracytoplasmic organelles but did not penetrate the nuclei (S1 Fig). Furthermore, neither probe was able to label plasma membrane as shown by experiments using CellMask™ Orange dye (S1A-E Figs) in larval tissues expressing mCD8-GFP (S1F-L Figs).

The intracellular distribution of NRB^MC009^ and NRB^ZLW0047^ was analyzed in more detail in *Drosophila* larval musculature in which we investigated the binding of these NRB fluorescent derivatives to ER, mitochondria, lysosomes and LDs. The larval body wall muscles provide a relatively simple system to study development of muscles, cytoskeleton dynamics, intracellular trafficking and neuromuscular junction dysfunction. In fact, besides the well-known actin and myosin filaments and their associated proteins, muscles also contain a cytoskeleton, intracellular organelles of the endo-lysosomal pathway, and well-defined endoplasmic reticulum and mitochondrial networks (25). We focused on these particular structures because we recently showed that they contained a significant density of binding sites for NRB^MC009^ (6). To study the intracellular localization of NRB^MC009^ we co-loaded it into larval muscle with organelle-specific red fluorescent dyes. To confirm NRB^ZLW0047^ localization we profiled it in larval muscles labelled with organelle-selective GFPs tagged proteins.

The results, showed good co-localization of NRB^MC009^ with ER tracker™ Red (ER probe, Pearson’s coefficient 0.65 ± 0.03, Fig 5A), Mitotracker™ Orange (mitochondrial probe, Pearson’s coefficient 0.54 ± 0.05, Fig 5B) and LipidTOX™ (lipid droplets probe, Pearson’s coefficient 0.42 ± 0.02, Fig 5C); however, NRB^MC009^ did not colocalize with Lysotracker™ Deep Red (a lysosome probe, Pearson’s coefficient 0.04 ± 0.04, Fig 5D). The localization of NRB^ZLW0047^ was confirmed using green fluorescent-tagged proteins that specifically targeted the ER (UAS-Hneu-GFP), mitochondria (UAS-Mito-GFP) and lysosomes (UAS-Lamp-GFP), and BODIPY 493/503 dye that targeted the LDs. The results, shown in Figs 5E-H, demonstrated that the distribution of NRB^ZLW0047^ partially overlapped that of NRB^MC009^; that both NRB^MC009^ and NRB^ZLW0047^ exhibited good labeling of ER and mitochondria (Pearson’s coefficient 0.51 ± 0.01 and 0.68 ± 0.01, respectively, Figs 5F and 5G); that both were absent in lysosomes (Pearson’s coefficient 0.04 ± 0.02, Fig 5I); and that they differed in their localization in LDs, where NRB^MC009^ fluorescence was present (Pearson’s coefficient 0.42 ± 0.02, see also Fig 5C) but NRB^ZLW0047^ fluorescence was absent (Pearson’s coefficient 0.16 ± 0.05, Fig 5H).

**Fig 5.**
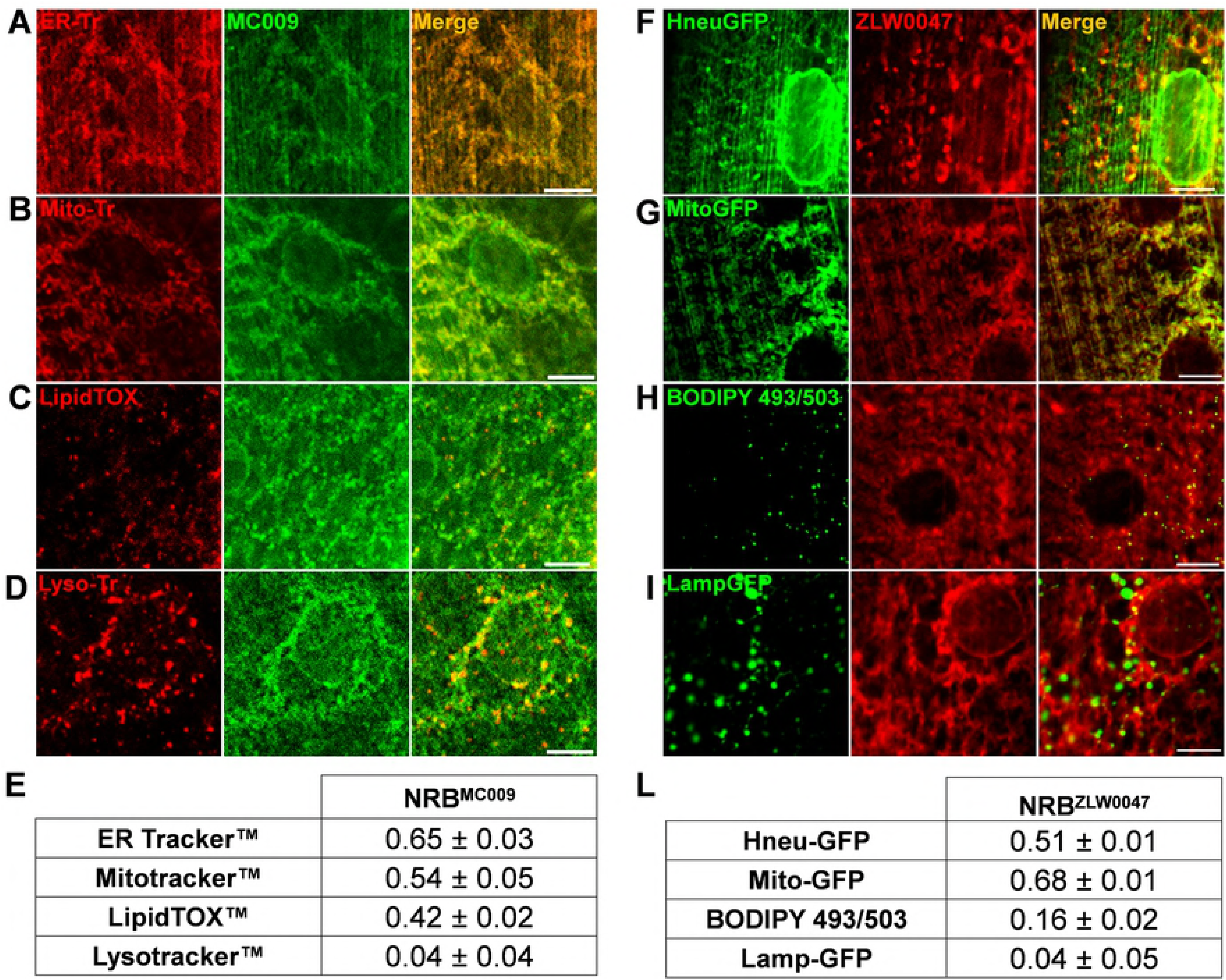
NRB^MC009^ and NRB^ZLW0047^ intracellular distribution in larval muscles. Confocal live imaging of w^[1118]^ third instar larval muscles 6-7 from segment A3 labeled and ER-tracker™ 2 µM (ER marker, **A**), Mitotracker™ 1 µM (mitochondria marker, **B**), LipidTOX™ 1:100 (LDs marker, **C**), and Lysotracker™ 2 µM (lysosomes marker, **D**), all in red, together with NRB^MC009^ 500 nM (green). Magnification 60x; scale bars 10 μm. Summary table of Pearson’s correlation coefficients between NRB^MC009^ and fluorescent organelle-marker probes used in larval muscles (**E**). Data are expressed as mean ± SEM, n≥10. Confocal live imaging of third instar larval muscles 6-7 from segment A3 of UAS-Hneu-GFP/Tubulin-Gal4 (ER marker, **F**), UAS-Mito-GFP/Tubulin-Gal4 (mitochondrial marker, **G**), w^[1118]^ added with BODIPY 493/503™ 10 μg/ml (LDs marker, **H**), and UAS-Lamp-GFP/+;Tubulin-Gal4/+ (Lysosomes marker, **I**), all labeled with NRB^ZLW0047^ 1 µM (red). Magnification 60x; scale bars 10 μm. Summary table of Pearson’s correlation coefficients between NRB^MC009^ and fluorescent organelle-markers in *Drosophila* larval muscles (**L**). Data are expressed as mean ± SEM, n≥10.

These data show that both compounds, but particularly NRB^MC009^, can be used to visualize and distinguish most organs/tissues of the dissected living larvae, and allow good definition of their intracellular structures. In addition, the co-localization studies revealed that both NRB^MC009^ and NRB^ZLW0047^ labelled subcellular organelles, that they preferentially targeted the ER and mitochondria, and that they were totally absent from the nuclei, plasma membranes and lysosomes, which is in agreement with data reported in mammalian cell studies (6). Moreover, the efficiency of the probes was tested with two different approaches: 1) in combination with commercially available dyes for live imaging, and 2) together with GFP tagged protein markers that bound to specific cellular structures. In both the experimental backgrounds the NRB based probes allowed the identification of endoplasmic reticulum and mitochondria structures, making both probes useful new markers in *Drosophila* studies. Based on the capability of these dyes to recognize the same intracellular structures in both mammalian and fruit fly cells, their potential use in more complex animal models is anticipated. Subsequently, our ongoing work is focused on the development of these probes as tools to allow live imaging studies to be conducted in mouse and rat tissues.

### NRB^MC009^ and NRB^ZLW0047^ cellular distribution in pathologic mutation-related phenotypes

In consideration of the preferential distribution of NRB-derived fluorescent probes to ER and mitochondria we next verified if they could be developed into tools to highlight phenotypic modifications of the ER and mitochondrial networks in *Drosophila* muscles. A large number of human disease genes are conserved in *Drosophila* and its genome can be easily manipulated to recreate and study human pathologic phenotypes (26); subsequently, Drosophila is widely used as a model to study muscle growth, degeneration and correlated diseases (Beckett and Baylies, 2006; Hirth, 2010; Kreipke et al., 2017; McGurk et al., 2015; Rossetto et al., 2011).

In this study NRB^MC009^ and NRB^ZLW0047^ were tested on two *Drosophila* pathologic models: Charcot– Marie–Tooth disease (CMTd) and hereditary spastic paraplegia (HSP) (31,32).

CMTd *Drosophila* phenotype was obtained by inducing a downregulation of *Marf*, the fruit fly orthologue of the human gene *Mitofusin2*, which encodes for a GTPase that, together with Opa1, fuses mitochondria; mutations of this gene are implicated in CMT disease (33). The depletion of this protein in *Drosophila* is known to cause fragmented and clustered mitochondria in neuronal cell bodies and to disorganize the typical sarcomeric location of mitochondria in the larval muscles, clumping them mainly around the nuclei (34). Fig 6B shows the fluorescent distribution of NRB^MC009^ and NRB^ZLW0047^ in muscles of larvae in which *Marf* had been ubiquitously downregulated (UAS-Marf^RNAi^/Tubulin-Gal4). Since these fluorescent images are comparable with those previously reported with the mitochondrial marker UAS-Mito-GFP (34) in the same model, when considering the mitochondrial labeling properties of NRB^MC009^ and NRB^ZLW0047^ (this study), it can be argued that fluorescent derivatives of NRB are able to also stain altered mitochondria, and be able to highlight pathologic mitochondrial phenotypes.

**Fig 6.**
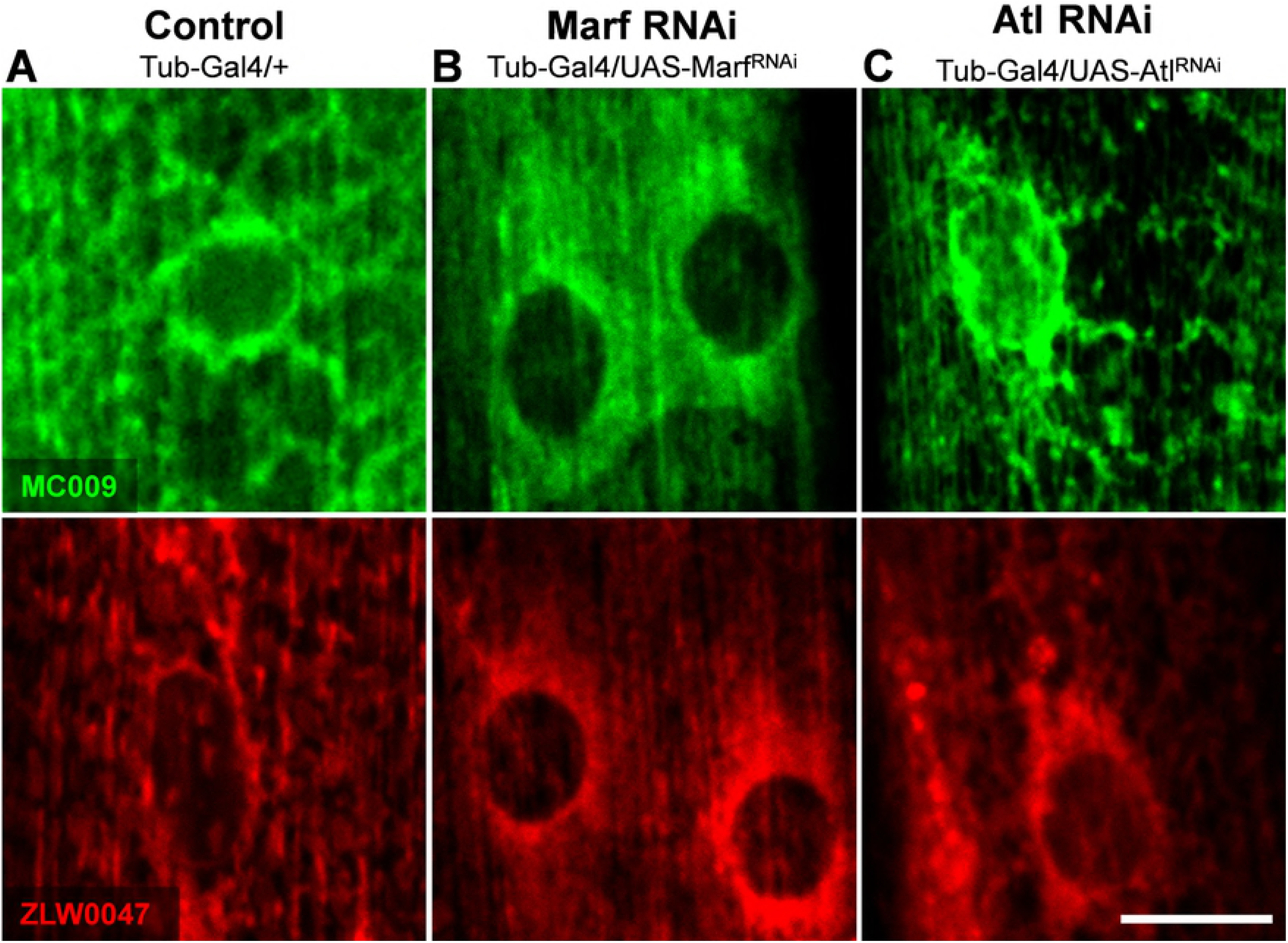
NRB^MC009^ and NRB^ZLW0047^ highlight pathologic mutation related phenotypes. Confocal live imaging of *Drosophila* muscles 6-7 from segment A3 of (**A**) control (Tubulin-Gal4/+), (**B**) Marf downregulation (UAS-Marf^RNAi^/Tubulin-Gal4), and (**C**) atlastin downregulation (UAS-Atlastin^RNAi^/Tubulin-Gal4) labeled with NRB^MC009^ 500 nM (green) or NRB^ZLW0047^ 1 µM (red). Magnification 60x; scale bars 10 μm.

HSP *Drosophila* phenotype was obtained by inducing a downregulation of atlastin. Atlastins are membrane-bound dynamin-like GTPases implicated in ER network morphogenesis, and mutations in *atlastin1* gene are involved in the onset of a common form of HSP (SPG3A). *Drosophila* holds a unique highly conserved *atlastin* orthologue, and its downregulation elicits a fragmented ER in neurons and an enrichment of ER punctae localized in the proximity of nuclei, and visualized using UAS-KDEL-GFP (18). Labeling of larva fillets with NRB^MC009^ and NRB^ZLW0047^ revealed a different pattern between muscles of wild type (Fig 6A) and atlastin-downregulated (UAS-Atl^RNAi^/Tubulin-Gal4) larvae (Fig 6C), in which a brighter perinuclear signal, compatible with the previously described HSP phenotype, is observed.

Taken together, these results indicate that NRB^MC009^ and NRB^ZLW0047^ could be useful tools for *Drosophila* live imaging to highlight phenotypes attributable to mutations in, and/or downregulation of genes implicated in mitochondria and/or endoplasmic reticulum network modifications. In addition, the short time it takes for these probes to permeate and label tissue and their general ease of use, means that both could be used as tools in compound screening studies to identify candidates that would help alleviate any network malfunction due to genetic modification.

### NRB^MC009^ and NRB^ZLW0047^ in food intake tests

Next we explored the possibility of using NRB^MC009^ and NRB^ZLW0047^ as tools to evaluate food intake and to investigate potential gut morphological modifications in *Drosophila in vivo*. By adopting a three-choice test (behavioral choice test) we verified that NRB^MC009^ and NRB^ZLW0047^, when added to the food, were accepted by the flies. As summarized in Fig 7A, there were no substantial differences in food preference between the standard diet and NRB^MC009^- and NRB^ZLW0047^-supplemented diets, indicating that the presence of the dyes did not influence the larval food choice. In addition, fluorescence imaging of larvae fed for 30 min with probe-enriched liquid food indicated that both NRB^MC009^ and NRB^ZLW0047^ could be clearly detected in the gut (Fig 7B); a more in depth analysis revealed that gut fluorescence was regulated by the probes contained in the food, since no signal was observed in the gut wall, leading to the conclusion that the strength of the fluorescent signal could be taken as an index of the quantity of ingested food. Fig 7C reports the results of the food intake assay, expressed as a percentage of the gut stained area relative to the total body area - no significant difference was observed between larvae fed with NRB^MC009^, NRB^ZLW0047^, or brilliant blue dye, a commonly used dye for the evaluation of food intake in *Drosophila* (21). The lack of gut labeling by NRB^MC009^ and NRB^ZLW0047^, although a useful outcome for the food intake test, was somewhat unexpected, particularly considering the results obtained in the dissected larvae (see Figs 2 and 3) where the dyes were clearly localized to the intestinal tract. To explain this inconsistency, we hypothesized that the time of exposure (30 min) of the larvae to the probe-supplemented food in the food intake assay was potentially too short to allow an internalization of the dyes to the gut epithelial cells. Therefore, we analyzed the gut wall of larvae grown for 5-7 days in food enriched with NRB^MC009^ or NRB^ZLW0047^. Figs 7D-F show the clear difference between vehicle-fed (Fig 7D) and NRB^MC009^- and NRB^ZLW0047^-fed larvae (Figs 7E and 7F, respectively). The bright fluorescent signal in the digestive tract (mainly midgut and hindgut) indicates that *Drosophila* larvae readily eat the probe-containing food. The digestive tract of NRB^MC009^- and NRB^ZLW0047^-fed larvae were clearly labeled by the dyes, a result that was accentuated by the absence of fluorescence in the rest of the body. The *Drosophila* intestinal tract is formed by a monolayer of epithelial cells, intestinal stem cells and enteroendocrine cells, surrounded by visceral muscles, nerves and tracheae. Ingested food from the proventriculus is pushed into the midgut, the main region of digestion and absorption, and then to the hindgut where the final absorption process takes place (13,14). A deeper investigation on dissected *Drosophila* gut revealed that NRB^MC009^ and NRB^ZLW0047^ did not only label the food that was present and visible in the intestinal tract (Figs 7I and 7L), but they also bound to the gut external muscular cells (Figs 7G and 7J) and enterocytes (Figs 7H and 7K).

**Fig 7.**
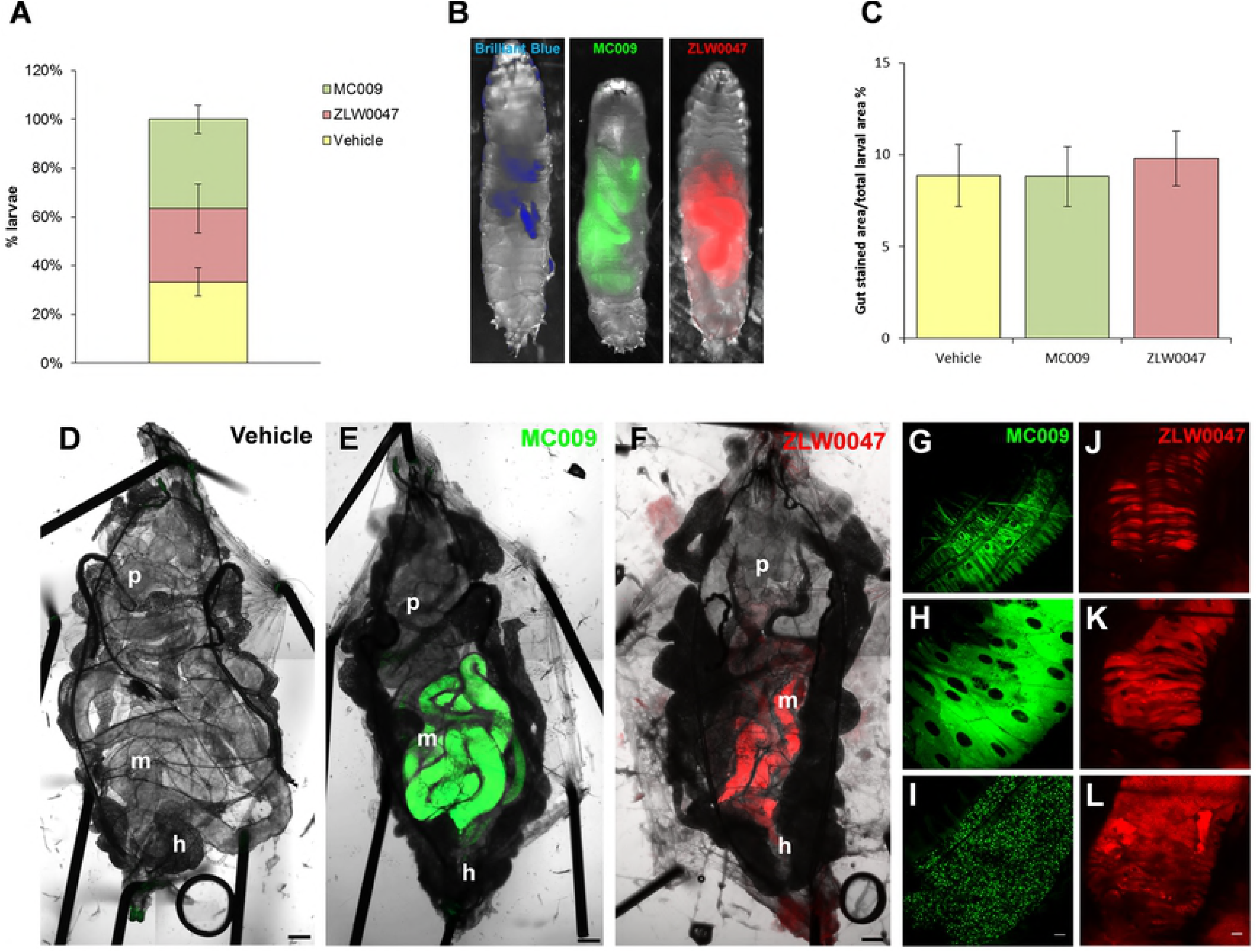
Use of NRB^MC009^ and NRB^ZLW0047^as dyes for food intake tests. Quantification of larval dispersal after 5 minutes of three-choice preference assay of vehicle-added food, NRB^MC009^-added food (20 µM), or NRB^ZLW0047^-added food (20 µM). Data were expressed as percent of total larvae number and represent mean ± SEM of five different experiments (**A**). Representative images of larvae fed with liquid food supplemented with Brilliant blue R 0.08%, NRB^MC009^ 20 µM, or NRB^ZLW0047^ 20 µM, where gut was labeled by the three dyes (**B**) and quantification of gut stained area versus total larval area (**C**). Data were expressed as mean of percent ± SEM of 30 larvae. Confocal live imaging of whole dissected *Drosophila* third instar larva fed with vehicle-supplemented food (**D**), NRB^MC009^ 20 µM-supplemented food (**E**), or NRB^ZLW0047^ 20 µM-supplemented food (**F**); magnification 5x; scale bar 200 μm. p: proventriculus, m: midgut, h: hindgut. Detailed images of mid gut external muscular cells (**G**, **J**), enterocytes (**H**, **K**) and intestinal food (**I**, **L**), labeled with the two NRB fluorescent derivatives; magnification 40x; scale bars 20 μm. The bright fluorescence of NRB^MC009^ and NRB^ZLW0047^ make these probes eminently suitable for use in food intake tests and chronic feeding assays; as monitoring tools for abnormal gut morphology; and identifying defects in gut functionality during development or screening tests.

### NRB^MC009^ and NRB^ZLW0047^ toxicity

In an attempt to validate the use of the NRB-derived fluorescent probes for use in chronic assays, we next verified their lack of toxicity in *Drosophila* by exposing the flies to NRB^MC009^ or NRB^ZLW0047^ over their entire life-cycle. The results indicate that male and female flies readily ingested NRB^MC009^- and NRB^ZLW0047^-supplemented food, mated and laid eggs normally, from which embryos hatched and larvae developed, grew, underwent pupation and eclosed in a similar fashion to non-treated flies. In addition, no difference in eclosion rate and lethality of adult flies was observed between probe-exposed and control flies (Fig 8A). Finally, we could not detect any apparent macroscopic morphological alteration in adult flies treated with either NRB^MC009^ or NRB^ZLW0047^ (Fig 8B).

**Fig 8.**
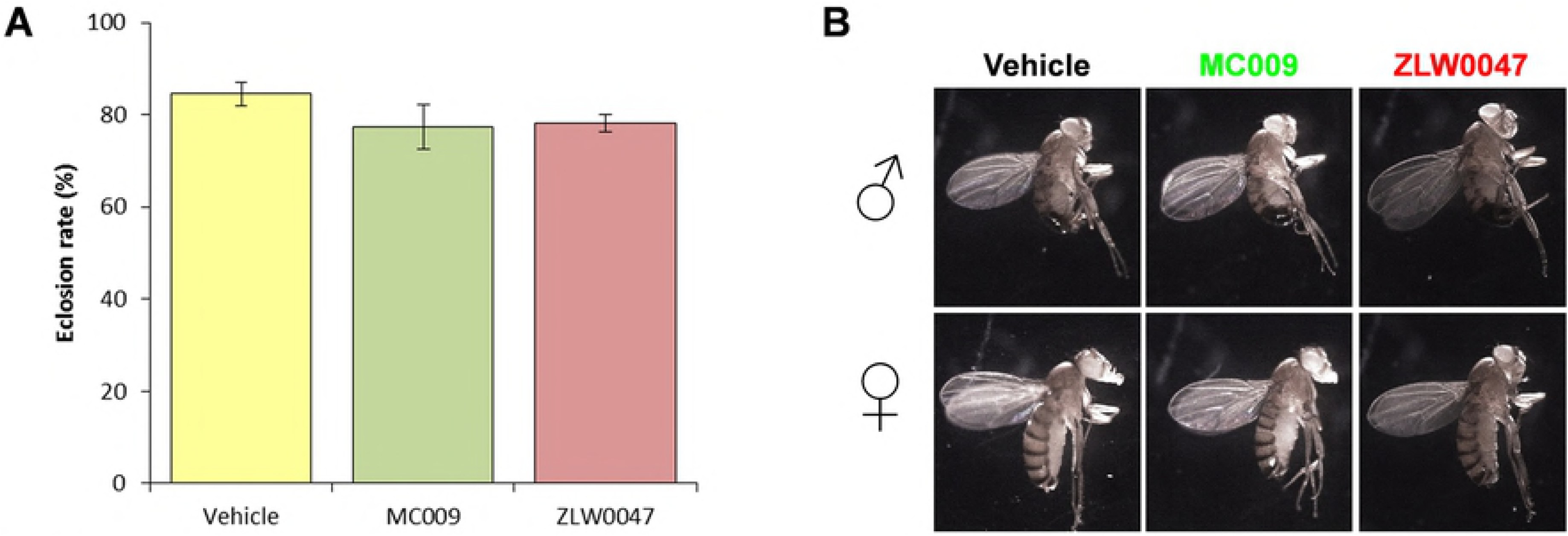
NRB^MC009^ and NRB^ZLW0047^ chronic toxicity. Evaluation of toxicity after exposition to standard food added with vehicle, NRB^MC009^ (20 µM) or NRB^ZLW0047^ (20 µM) over the *Drosophila* entire life-cycle. (**A**) Quantification of eclosion rate. Data were expressed as mean of percent of emerged flies versus to the total number of pupae. (**B**) Representative images of emerged male (♂) and female (♀) w^[1118]^ flies, where morphological alterations of body, eyes, wings, and legs were evaluated.

The absence of toxicity and the high level of palatability supportsNRB^MC009^ and NRB^ZLW0047^ as potential monitoring tools for long term feeding assays; as markers of intestinal epithelia; and their use in studying *Drosophila* digestive tract functionality e.g. monitoring the effect of compounds or diet on intestinal performance (Apidianakis and Rahme, 2011; Gasque et al., 2013; Storelli et al., 2018)These attributes also make these probes potentially useful as mammalian gastrointestinal (GI) tract markers; for example, the GI tract is one of the most studied tissues in many pathological rodent models (38,39) and the availability of easy-to-use, high performance fluorescent probes to detect intracellular structure abnormalities could be of great benefit.

Overall, this study has investigated the imaging applications of NRB^MC009^ and NRB^ZLW0047^ in *Drosophila melanogaster* in vivo. The analysis of the fluorescent signals of these compounds reveal that both can label subcellular specific organelles (preferentially ER and mitochondria), in both wild type and pathological phenotypes. The absence of toxicity and the minimal effect on palatability also allows them to be used as potential monitoring tools in feeding assays, and as markers for intestinal epithelia that could be useful in *Drosophila* digestive tract studies. In summary, the characteristically bright signals of NRB^MC009^ and NRB^ZLW0047^, in combination with their capacity to permeate tissues rapidly, makes them eminently suitable for confocal imaging applications. Our future studies will focus on investigating whether these compounds, with their enhanced attributes, may have potential to be used in invertebrate animal models.

## Supporting information

**S1 Fig. NRB^MC009^ and NRB^ZLW0047^ do not label plasma membranes nor nuclei in larval tissues.**

Confocal live imaging of w^[1118]^ larval (A) peripodal membrane cells of a leg imaginal disc, (B) salivary gland, (C) fat body, and (D) central nervous system labeled with NRB^MC009^ 500 nM (green) and CellMask™ Orange (cell membrane marker) 1 µM (red). Magnification 60x, scale bars 10 μm (A-C); magnification 40x, scale bar 100 μm (D). N: nucleus, G: ganglion, Nv: nerves. Summary table of Pearson’s correlation coefficients between NRB^MC009^ and CellMask™ in the evaluated tissues (E). Data expressed as mean ± SEM, n≥10. Confocal live imaging of UAS-mCD8-GFP/Tubulin-Gal4 (cell membrane marker) larval (F) peripodal membrane cells of a leg imaginal disc, (G) salivary gland, (H) fat body, and (I) central nervous system labeled with NRB^ZLW0047^ 1 µM (red).

Magnification 60x, scale bars 10 μm (A-C); magnification 40x, scale bar 100 μm (D). N: nucleus, G: ganglion, Nv: nerves. Summary table of Pearson’s correlation coefficients between NRB^ZLW0047^ and mCD8-GFP signal in the evaluated tissues (L). Data expressed as mean ± SEM, n≥10.

